# *Shewanella oneidensis* grows on biochar as an electron acceptor via direct contact without flavins

**DOI:** 10.64898/2026.09.21.753187

**Authors:** Ankit Singh, Jessica L. Keffer, Jiwon Choi, Pei C. Chiu, Clara S. Chan

**Author notes:** Corresponding authors: Jessica L. Keffer and Clara S. Chan Department of Earth Sciences, University of Delaware, Newark DE 19716.

## Abstract

Pyrogenic black carbon, such as biochar and wildfire char, possesses a reversible electron storage capacity (ESC), allowing it to serve as an electron donor or acceptor for microorganisms. However, not all the ESC is bioavailable, and the proportion available varies by culture. We hypothesized microbes capable of producing soluble redox mediators could access the ESC in biochar interior and thus utilize a higher percentage of the ESC than organisms without a mediator. To test this hypothesis, we grew wild-type *Shewanella oneidensis* (MR-1) and Δ*bfe* MR-1, a mutant incapable of exporting flavins, on air-oxidized wood biochar. Cell counts and redox titration show that both strains of MR-1 grew to similar cell densities and accessed comparable amounts of biochar’s ESC (∼19%). The addition of exogenous flavin mononucleotide (FMN) yielded no measurable increase in final cell density, due to the strong adsorption of FMN to biochar. SEM images show attachment of MR-1 and Δ*bfe* to biochar surfaces and the presence of nanowires. This study demonstrates MR-1 can utilize biochar’s ESC for growth, through direct physical contact without the involvement of flavins. Understanding how microbes metabolize biochar provides insight into the impacts and biogeochemical roles of pyrogenic carbon.

## Introduction

Biochar is a subset of black carbon derived through the pyrolysis of organic material. Black carbon is an integral part of the global carbon cycle (produced naturally at >10^14^ g per year) and the biochar subset is the basis for a growing industry through its use as an amendment in agricultural soils (Coppola *et al*. 2018, Liu *et al*. 2023, Fortune Bus. Insights 2026). Biochar derived from plant biomass is redox-active and has a constant, reversible electron storage capacity (ESC), which allows it to accept and/or donate electrons (Xin, Xian, and Chiu 2019). The fact that ESC is a property of all plant-derived biochar means it has the capability to play a role in microbial respiration pathways and chemical redox processes in natural and engineered systems (Kappler *et al*. 2014, Saquing, Yu, and Chiu 2016, Xin *et al*. 2023, Choi, Xin, and Chiu 2025). However, this capability has not been explored well, so it is not clear how the redox properties of biochar impact soil microbial communities and function.

Previous studies have demonstrated the capacity of different microorganisms to interact with and grow using biochar (Saquing, Yu, and Chiu 2016, Yu *et al*. 2016, Xin *et al*. 2023, Li W *et al*. 2025). Biochar can be used by *Geobacter metallireducens* GS-15 for acetate oxidation and nitrate reduction (Saquing, Yu, and Chiu 2016), and we have shown it can be a terminal electron acceptor in *G. metallireducens* GS-15 growth with acetate and fully oxidized biochar (Li W *et al*. 2025). *G. metallireducens* GS-15 could use ∼20% of the ESC of oxidized biochar (Li W *et al*. 2025), which likely represents the surface-available portion of the ESC. However, in another study, a wastewater mixed community culture accessed ∼50% of the ESC (Xin *et al*. 2023), suggesting different microbes interact with biochar in different ways. Indeed, several mechanisms of microbial extracellular electron transfer (EET) with biochar have been postulated including direct contact, electron shuttling, and the production of extracellular polymeric species (Chen S *et al*. 2014, Kappler *et al*. 2014, Yang Z *et al*. 2022, Li W *et al*. 2025). Extracellular electron shuttles, including microbially-produced flavins, phenazine, quinones, and environmental AQDS and dissolved organic matter, have been shown to enhance microbial EET with other solid substrates, such as Fe(III)-oxides and electrodes (Von Canstein *et al*. 2008, Kotloski and Gralnick 2013, Chen Z *et al*. 2017, Tolar, Li, and Ajo-Franklin 2023, Qin *et al*. 2024, Li J *et al*. 2026). However, it is unclear if electron shuttles are involved in microbial EET with biochar.

To further explore how microbes interact with biochar, we investigated *Shewanella oneidensis* MR-1 (hereafter MR-1). MR-1 is a facultative anaerobe with high metabolic versatility through its ability to respire substrates such as oxygen, organic compounds, inorganic ions, soluble metals, and various solids like metal oxides and electrodes (Myers and Nealson 1988, Tiedje 2002, Bernardet and Bowman 2006, Hunt *et al*. 2010, Kouzuma *et al*. 2015). MR-1 has the capability to perform EET through direct contact between cytochromes on the cellular surface and substrate or the formation of cytochrome-containing nanowires, and producing three different types of soluble electron shuttles: flavin mononucleotide (FMN), riboflavin, and flavin adenine dinucleotide (FAD) (Von Canstein *et al*. 2008, Jiang *et al*. 2010, Kotloski and Gralnick 2013, Barchinger *et al*. 2016, Sun *et al*. 2021). Flavins produced by MR-1 have dual roles, acting as intracellular redox cofactors for membrane-bound cytochromes and as extracellular electron shuttles (Kotloski and Gralnick 2013, Hong and Pachter 2016). While FMN is the primary flavin secreted during growth (Von Canstein *et al*. 2008), all flavins are exported through the same exporter protein Bfe (SO_0702) (Kotloski and Gralnick 2013). Flavins help MR-1 interact with poorly soluble electron acceptors (Von Canstein *et al*. 2008), as a MR-1 mutant with deletion of the flavin exporter gene produces less Fe(II) when provided with the insoluble iron oxyhydroxide ferrihydrite as the sole electron acceptor (Kotloski and Gralnick 2013). Since biochar is a solid electron acceptor, we hypothesize secreted flavins could allow MR-1 to access a larger percentage of biochar’s ESC compared to *G. metallireducens* GS-15.

In this study, we evaluated the ability of MR-1 to grow using biochar as its sole electron acceptor, with and without flavins. To determine whether extracellular flavins played a role, we observed the growth of three different treatments: wild-type MR-1, wild-type MR-1 with exogenous FMN, and a mutant MR-1 (MR-1 Δ*bfe*) without the flavin exporter gene (Kotloski and Gralnick 2013). We determined MR-1 growth on biochar was largely not influenced by flavins, regardless of whether the flavins were produced endogenously or added exogenously. We also observed contact between MR-1 and biochar through fluorescent and scanning electron microscopy and we also measured electron content of biochar samples through chemical redox titration with titanium(III) citrate. Through these analyses we provide insight into the flavin-independent growth and EET by MR-1 when using oxidized biochar as the sole electron acceptor.

## Materials and Methods

### Biochar

Soil Reef biochar (SRB), a commercial biochar produced through pyrolysis of Southern Yellow Pine wood chips at 550 °C (Saquing, Yu, and Chiu 2016), was sieved to 250–500 μm. To completely remove residual electrons in SRB and hence maximize its ESC, SRB was oxidized with dissolved O_2_ in continuously aerated deionized water for 10 days (Xin *et al*. 2023). Air-oxidized SRB was then vacuum-filtered, dried, autoclaved, and stored in a desiccator. Any residual O_2_ associated with SRB (either gaseous in pores or sorbed onto surfaces) was removed by vacuum for 24 h followed by equilibration with the anaerobic chamber atmosphere (∼97.5% N_2_, ∼2.5% H_2_) for 3 days before use.

### Cell cultures and counting

*Shewanella oneidensis* MR-1 and Δ*bfe* were a gift from Dr. Jeffrey Gralnick (University of Minnesota) (Kotloski and Gralnick 2013). MR-1 was cultured on a minimal *Shewanella* basal media containing (per liter) 0.46 g of NH_4_Cl, 0.225 g of K_2_HPO_4_, 0.225 g of KH_2_PO_4_, 0.117 g of MgSO_4_·7H_2_O, and 0.225 g of (NH4)_2_SO_4_. Prior to autoclaving, 5 ml of a mineral mix (containing per liter 1.5 g of nitrilotriacetic acid, 0.1 g of MnCl_2_·4H_2_O, 0.3 g of FeSO_4_·7H_2_O, 0.17 g of CoCl_2_·6H_2_O, 0.1 g of ZnCl_2_, 0.04 g of CuSO_4_·5H_2_O, 0.005 g of AlK(SO_4_)_2_·12H_2_O, 0.005 g of H_3_BO_3_, 0.09 g of Na_2_MoO_4_, 0.12 g of NiCl_2_, 0.02 g of NaWO_4_·2H_2_O, and 0.10 g of Na_2_SeO_4_) as well as 5 ml of a vitamin mix (containing per liter 0.002 g of biotin, 0.002 g of folic acid, 0.02 g of pyridoxine HCl, 0.005 of thiamine, 0.005 g of nicotinic acid, 0.005 g of pantothenic acid, 0.0001 g of B-12, 0.005 g of *p*-aminobenzoic acid, and 0.005 g of thioctic acid) was added (Baron *et al*. 2009). HEPES buffer (50 mM) was added, and media was buffered in the pH range 6.9 – 7.1. After autoclaving, 0.05% w/v of casamino acids were added. The media was bubbled with N_2_ gas for 1 hour and left in an anaerobic chamber for 24 hours to equilibrate to ensure deoxygenation prior to inoculation. Anaerobic cultures of wildtype (WT) and Δ*bfe* MR-1 were grown in the minimal media with 15 mM lactate and 20 mM fumarate. There was no observed difference in anaerobic growth between WT and Δ*bfe* on lactate/fumarate (**Fig. S1**). These lactate/fumarate cultures were washed and resuspended in media with lactate only and stored as glycerol stocks for inoculation of additional experiments. Experimental cultures contained 20 mM L-lactate and 1 g of oxidized SRB in 100 mL vials with 50 mL of media and 50 mL of headspace. One set of cultures was amended with 10 µM flavin mononucleotide monosodium dihydrate (ICN Biomedicals).

Samples of 0.5 - 1.5 mL of cultures were taken for cell counting, lactate, and flavin analysis. Each sample was vortexed, then after the biochar settled, samples of the liquid media were taken. Cell counting was performed by staining the cells with Syto 13 dye and counting on a Petroff-Hausser chamber using a Zeiss Axioimager Z1 fluorescent microscope. The rest of the sample portions were filtered through a 0.22 µM nylon syringe filter to remove cells and biochar and used for flavin and lactate analysis.

### Flavin analysis

Samples from the 10 µM FMN standard, cells + 10 µm FMN, cells + 10 µM FMN + 1 g biochar, 10 µM FMN + 1 g biochar, and media + 1 g biochar were filtered as above, then added to a UV-clear cuvette (BrandTech) and absorbance (200 - 600 nm) measured by an Agilent Cary 3500 UV-Vis spectrophotometer.

### Lactate assay

L-lactate was quantified using an enantiomer-specific enzymatic assay (Eton Bio). All samples were diluted 10x using DI water to fall within the standard curve. Fifty microliters of samples and 50 µL of enzyme solution were added to a 96 well plate and left to incubate at 37 °C for 30 minutes. Absorbance at 490 nm was measured using a plate reader (Molecular Devices Spectramax i3x). A standard curve was made using 0 - 3 mM sodium L-lactate (Sigma-Aldrich).

### ESC measurement

Biochar was recovered from cell culture vials after 14 days for ESC measurement through redox-titration with titanium (III) citrate. All filtering and washing steps were done in anaerobic conditions using deoxygenated DI water. Cultures were vacuum filtered through 47 mm glass microfiber filters. The collected biochar (∼1 g) was then added to conical tubes containing 50 mL of 1% Tween 80 solution (Xin *et al*. 2023) and shaken at 300 rpm for 45 minutes to remove cells and media residue. The washed biochar was then vacuum filtered on 47 mm glass microfiber filters and rinsed 3x with 1 L deoxygenated DI water to remove residual Tween 80. The triplicates were combined (∼3 g), and vacuum dried for 6 hours followed by open drying in the anaerobic chamber for 6 days. The ESC of air-oxidized and deoxygenated biochar from the stock used in these experiments and biochar retrieved at the end of the growth experiments were measured by redox titration using Ti(III) citrate as a reductant (− 0.36 V vs. SHE at pH 6.4) (Xin, Xian, and Chiu 2019). ESC was determined based on cumulative consumption of Ti(III) by the biochar, as measured by UV−Vis absorption at 400 nm. Since the electron donating capacity of air-oxidized biochar is zero, the measured ESC was taken to be the total ESC.

### Scanning electron microscopy (SEM)

Wild-type MR-1 and Δ*bfe* were grown as above, but with millimeter-sized pieces of unsieved oxidized biochar so that the biochar could be recovered for processing without disturbing attachment points between the cells and biochar. After 14 days growth, pieces of oxidized biochar were gently removed from the serum bottles with tweezers and fixed in 4% glutaraldehyde, washed three times, then dehydrated in an ethanol dilution series (25%, 50%, 75%, 95%, and 100%). The dehydrated samples were placed into a critical point dryer (Tousimis Autosamdri-815B), then mounted on Al stubs and coated with Pt on a Leica EM ACE600. Imaging was performed at 2kV on a Thermo Scientific Apreo Volumescope SEM.

### Results and Discussion

We tested the ability of wild-type MR-1 to grow while using biochar as a terminal electron acceptor. MR-1 grew with 20 mM lactate as the sole electron donor and 1 g of biochar as the sole electron acceptor in minimal media (**Fig. 1**). Cell density increased 60-fold to 2.93 (±0.59)*10^8^ cells/mL, reaching stationary phase within five days. Without biochar, in contrast, no growth was observed over the same period. We also grew MR-1 Δ*bfe*, which lacks the efflux transporter encoded by SO_0702, previously shown to be responsible for the export of extracellular flavins (Kotloski and Gralnick 2013). The Δ*bfe* mutant grew 58-fold to 2.77 (±0.63)*10^8^ cells/mL (**Fig. 1**). The similar growth curves and final cell densities of the two strains suggests that either flavins do not improve the utilization of biochar as an electron acceptor by MR-1, or that the wild-type MR-1 was not actively producing and/or exporting extracellular flavins, which is a possibility since the low levels of naturally secreted flavins by MR-1 have been shown to limit efficient EET in previous studies (Yang Y *et al*. 2015).

**Figure 1.**
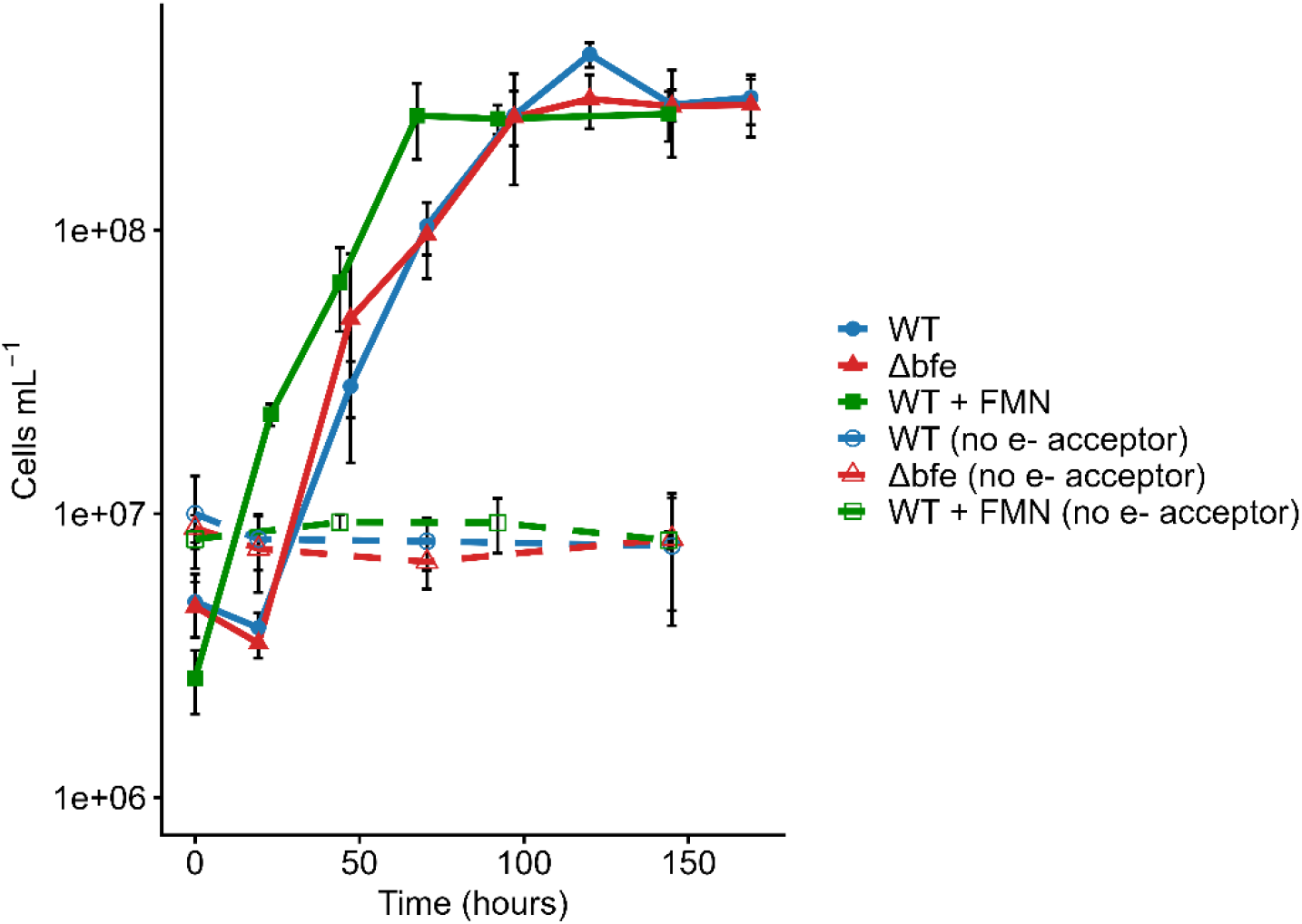
Growth of wild-type MR-1 (WT; blue circles), MR-1 Δ*bfe* (red triangles), and WT + 10 µM FMN (green squares) using biochar as sole terminal electron acceptor, compared to WT (open circles), Δ*bfe* (open triangles), and WT + 10 µM FMN (open squares) without an electron acceptor. Error bars represent one standard deviation from three replicates.

To test whether higher concentrations of FMN could cause growth differences, we grew wild-type MR-1 with 10 µM exogenous flavin mononucleotide (FMN) added in the same minimal media with either 1 g of biochar or no electron acceptor. The wild type with exogenous FMN grew 97-fold to 2.57 (±0.51)*10^8^ cells/mL (**Fig. 1**). These results show that the presence of exogenous FMN had no effect on the maximum growth yield of MR-1 on biochar. However, there was an observed difference in the time (∼24 hrs) that the FMN-amended MR-1 began exponential growth and stationary phase. Both the wild type without exogenous FMN and the Δ*bfe* mutant entered the exponential growth phase between 24-40 hours after inoculation while the wild type with exogenous FMN entered it prior to the first cell count at 24 hours (**Fig. 1**). The wild type with exogenous FMN also exited the exponential phase ∼24 hrs before the other two treatments. These results indicate that the presence of extracellular FMN in the system may allow MR-1 to decrease its lag phase and enter and exit growth earlier compared to when FMN is not already present. Because FMN is a cofactor for outer membrane cytochromes and can also be readily converted into riboflavin, the redox cofactor of OmcA (Okamoto *et al*. 2014, Hong and Pachter 2016), it is possible that the exogenous FMN reduced cellular FMN production requirements, expediting initial growth. However, the extent of growth and growth rate are similar between all three conditions suggesting FMN does not play a role as an extracellular electron shuttle in MR-1’s ability to access biochar’s ESC.

We noted an initial discrepancy at time 0 between the cell counts in the liquid phase of treatments with biochar compared to treatments without biochar, despite starting with identical inoculant. The number of WT MR-1 cells initially counted in the biochar treatment was around two-fold less than the no biochar control (4.9 (± 1.2)*10^6^ cells/mL and 1.0 (± 0.36)*10^7^ cells/mL, respectively; **Fig. 1**). The number of Δ*bfe* mutant cells initially counted in the biochar treatment was also around two-fold less than the control (4.7 (± 1.0)*10^6^ cells/mL and 8.9 (± 0.98)*10^6^ cells/mL, respectively; **Fig. 1**). The number of WT MR-1 cells initially counted in the biochar with 10 µM FMN treatment was around three-fold less than the control (2.6 (± 0.67)*10^6^ cells/mL and 8.1 (± 0.61)*10^6^ cells/mL, respectively; **Fig. 1**). The time 0 cell counts were taken <30 minutes after inoculation, indicating that initial association between MR-1 and biochar happened quickly, and was stable to physical vortexing, as cells were clearly visible remaining on the biochar particles after preparation for cell counting (**Fig. 2**). This likely leads to an underestimate of total cell counts, since we count only planktonic cells, although all conditions would be similarly undercounted.

**Figure 2.**
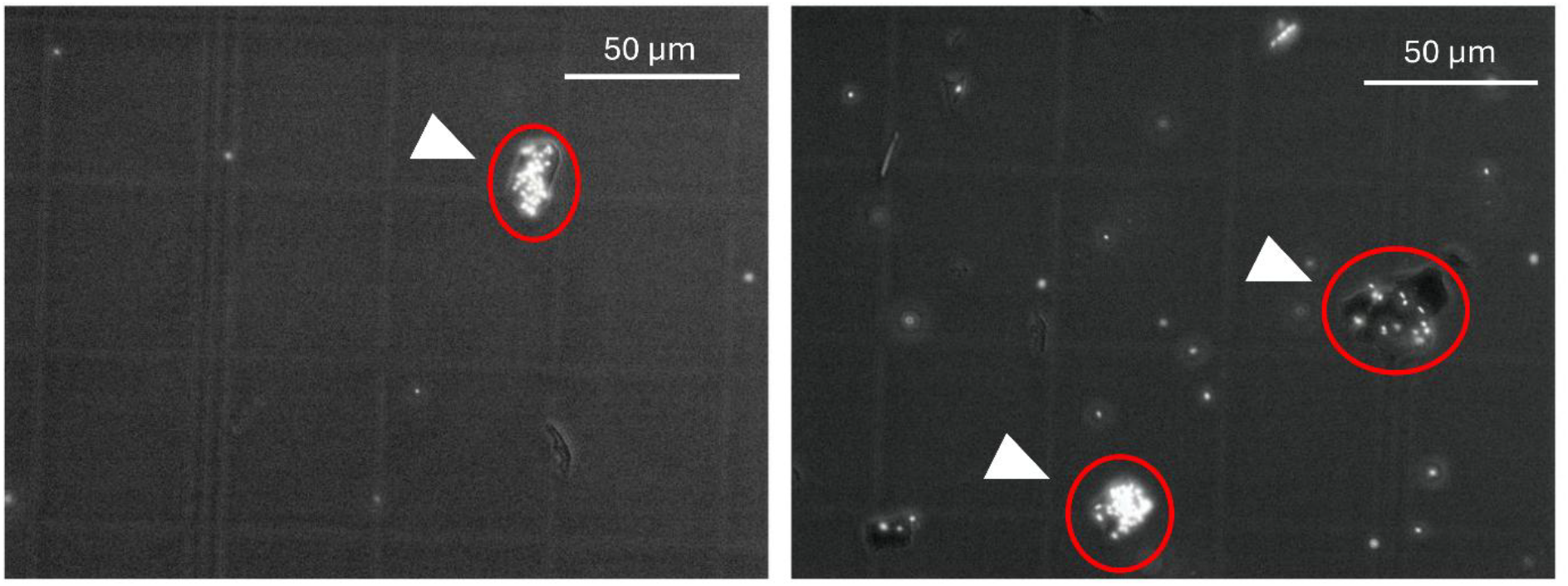
Fluorescent images of Syto 13-stained WT MR-1 (left panel) and Δ*bfe* (right panel) showing a high proportion of cells associated with biochar compared to the liquid media. Arrows indicate biochar particles, circled in red. The grid cells of the Petroff Hausser counting chamber are visible, which are 50 microns.

Lactate measurements show that consumption of lactate corresponded to MR-1 growth on biochar for both strains, whereas minimal lactate was consumed without biochar (**Table S1**). Given that each lactate molecule can donate four electrons (Eq. 1), the total e^-^ derived from lactate based on net lactate utilization was 0.99 mmol for the wild type and 1.2 mmol for the Δ*bfe* strain in 50 mL cultures (**Table 1**). These results further indicate that the ability to secrete flavins had no effect on MR-1 growth on biochar.

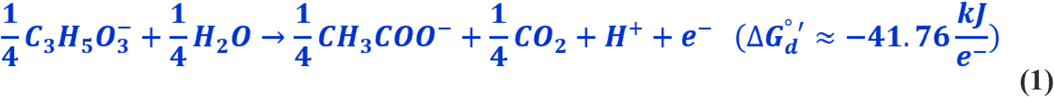

**Table 1.** Net electron consumption and donation, and total percentage of biochar ESC used.

| Strain | Net $e^-$ consumed<sup>a</sup> | Net $e^-$ deposited<sup>b</sup> | % of biochar ESC used |
| --- | --- | --- | --- |
| WT | 0.99±0.76 | 0.52±0.11 | 18±4.2 |
| $\Delta bfe$ | 1.2±0.33 | 0.66±0.05 | 20±1.8 |

<sup>a</sup>Calculated from electron donor (lactate) measurements
<sup>b</sup>Calculated from electron acceptor (biochar) measurements

The electron storage capacity measured for this batch of biochar was 3.4±0.04 mmol/g (**Table S1; Fig. S2**). The total number of electrons that wild-type MR-1 (0.52±0.11 mmol) and Δ*bfe* mutant (0.66±0.05 mmol) deposited into oxidized biochar during growth were not significantly different (**Table 1**). These values represent ∼18-20% of the measured total ESC. Due to the presence of excess electron donating capacity from lactate (since only 6-7 mM of the 20 mM provided was consumed, **Table S1**), it can be assumed that stationary phase was reached due to the depletion of bioavailable biochar ESC. The calculated electron consumption from lactate and direct measurements of biochar ESC support the conclusion that the wild type and mutant cultures are able to utilize a similar amount of the biochar’s ESC. This range is comparable to that reported for *G. metallireducen*s GS-15, which was able to access ∼19% of biochar’s total ESC (Li W *et al*. 2025). Because endogenous flavins do not appear to be increasing the ESC that is accessible to MR-1, and *G. metallireducens* GS-15 has been previously shown to not produce soluble electron shuttles (Nevin and Lovley 2000, Afkar *et al*. 2005), we conclude that the amount of ESC accessed by both of these strains (18-20%) represents the surface biochar ESC that is physically accessible to direct contact with a cell.

In order to determine whether there were any abiotic interactions occurring between FMN and biochar that could explain its lack of effect on MR-1’s maximum growth capacity, we monitored the interaction between FMN and biochar via absorbance spectroscopy. Oxidized FMN displays maximum absorbance between 400-500 nm (Macheroux 1999). Spectra from FMN in the liquid phase with no biochar show a clear absorption peak at ∼460 nm whether or not MR-1 cells are present, while treatments with FMN and biochar present together show no peak at ∼460 nm, suggesting that FMN was removed from solution by biochar via adsorption (**Fig. 3**). The samples were measured within 5 minutes of adding all substrates into the media, meaning the sorption of FMN to biochar happens relatively quickly, removing bioavailable FMN from the growth media. The large surface area of biochar and its variety of surface functional groups has been previously shown to adsorb organic molecules of various sizes and structures (Peng *et al*. 2016, Xing *et al*. 2023, Loebsack *et al*. 2025). The removal of FMN from the liquid media by biochar sorption is consistent with the cell count and electron consumption data, indicating that flavins do not enhance the capacity of MR-1 growth on biochar through electron shuttling.

**Figure 3.**
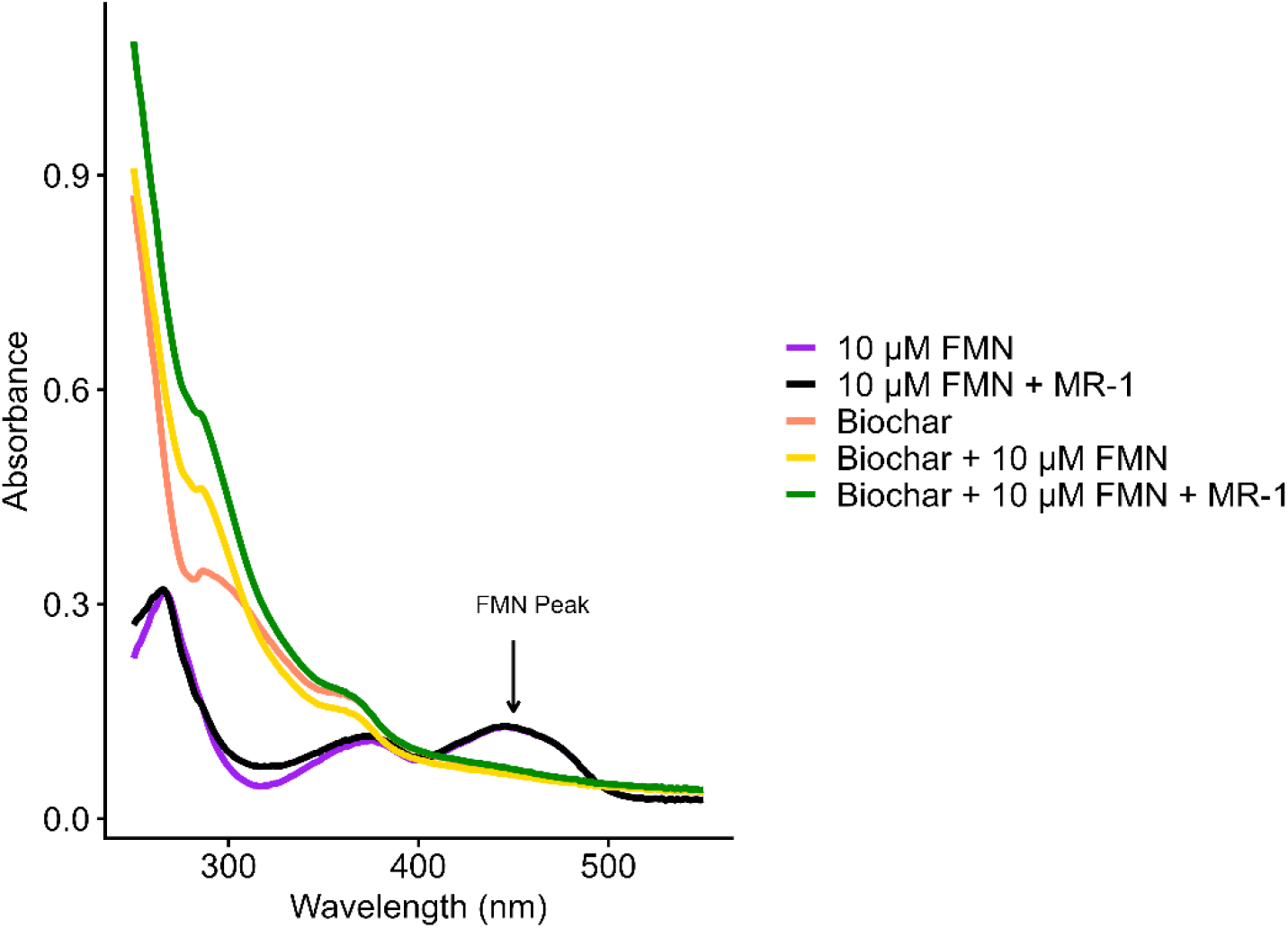
Absorbance measurements of five treatments: a 10 µM FMN standard (purple), media + 1 g biochar (orange), media + 1 g biochar + 10 µM FMN (yellow), media + wild-type MR-1 + 10 µM FMN (black), and media + 1 g biochar + wild-type MR-1 + 10 µM FMN (green). The characteristic FMN absorbance peak is indicated at approximately 450 nm.

MR-1 and *G. metallireducens* GS-15 both access a similar percentage (∼19%) of the total ESC of biochar (**Table 1** and (Li W *et al*. 2025)), which is likely the surface-available electron storage sites that can be utilized via direct contact. While FMN does not promote access to additional internal electron storage sites, it is possible that other redox-active molecules with different physiochemical properties would continue to function as soluble shuttles in the presence of biochar. Thus, organisms and environments with a more varied repertoire of electron shuttles that do not sorb to biochar, or communities producing an excess of shuttles that would allow some to be bioavailable even if others are sorbed, could use a higher percentage of biochar ESC like the previously observed wastewater community (Xin *et al*. 2023).

To further explore the interaction between biochar and MR-1, we prepared cultures for SEM using large pieces of biochar that would preserve cell-biochar spatial relationships with minimal disturbance during sample processing. We observed prevalent attachments between the cells and biochar, and both MR-1 and Δ*bfe* were similarly attached (**Fig. 4; Fig. S3**). Observed attachments frequently resembled smooth filaments near the cell and terminated in vesicle chains. There was no noticeable difference between MR-1 and Δ*bfe* in terms of the number or types of connections. The size and morphology of these attachments matches those previously shown to be nanowires composed of outer membrane extensions localized with cytochromes in MR-1 (Pirbadian *et al*. 2014, Subramanian *et al*. 2018). Thus, it is likely that MR-1 is utilizing direct attachment of its membranes and cytochromes to access the surface ESC of biochar to support its growth.

**Figure 4.**
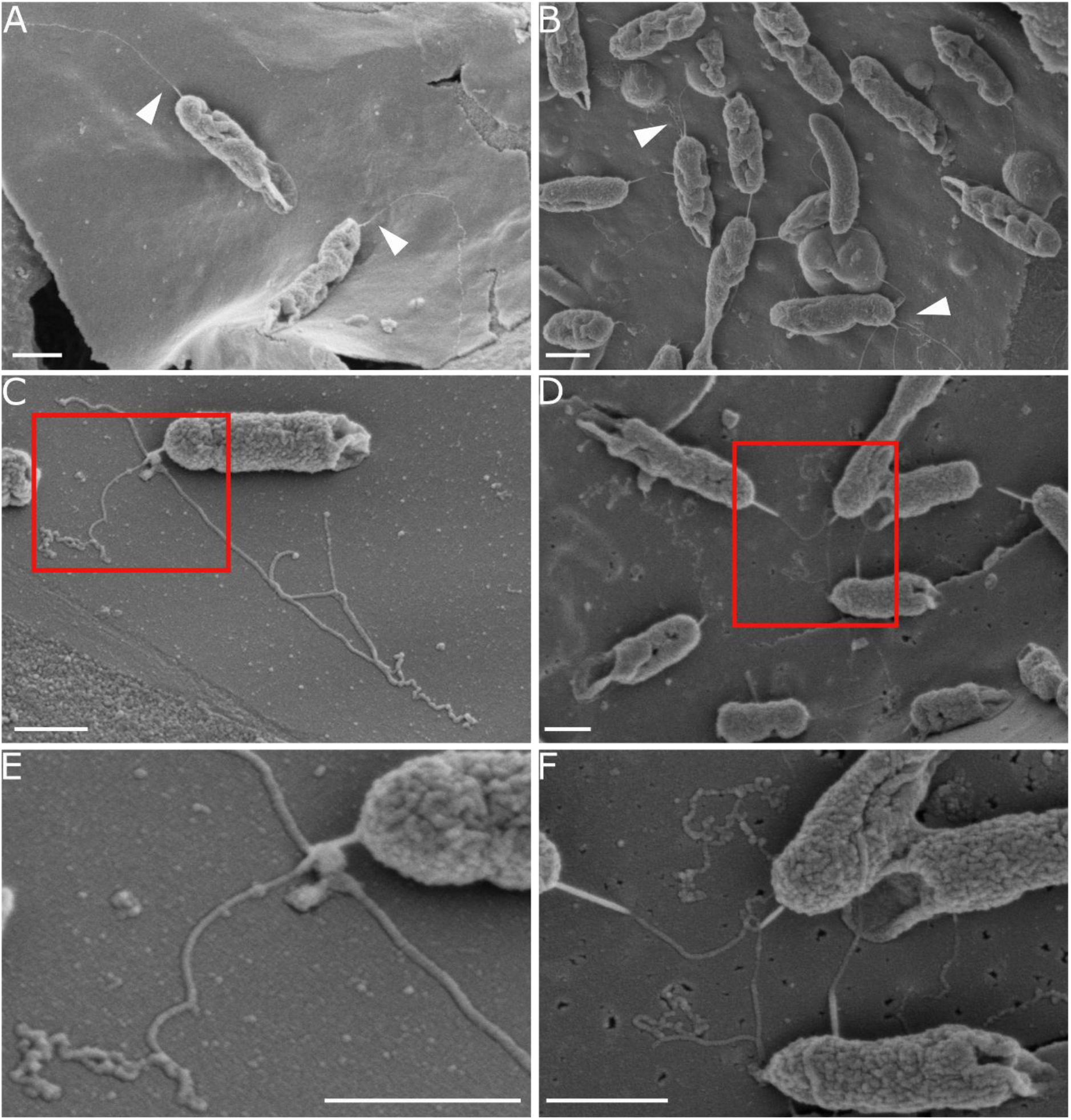
SEM images of wild-type MR-1 (A, C, E) and Δ*bfe* (B, D, F). White arrowheads in A and B show some examples of the nanowires. Red boxes in C and D are enlarged in E and F. All scale bars are 0.5 µm.

## Conclusions

This work represents a clear demonstration of growth of *S. oneidensis* MR-1 using oxidized biochar as the sole terminal electron acceptor (**Fig. 1**), adding to the list of substrates that MR-1 can use for EET. We observed that MR-1 accesses the electrons of biochar via direct contact, as many cells were seen firmly attached to biochar (**Fig. 2**) and it appears to be producing nanowires when growing on the biochar (**Fig. 4; Fig. S3**). In further support of direct contact as the mechanism for electron transfer with biochar, we observed that flavins do not appear to promote growth in MR-1 (**Fig. 1**) and FMN, the main flavin produced by MR-1 during growth, sorbs to biochar (**Fig. 3**), making it unavailable for redox shuttling. MR-1 has been reported to make these nanowires under conditions of electron acceptor limitation (Gorby *et al*. 2006), which agrees with our findings here where lactate is not fully consumed (**Table S1**) and MR-1 is limited to the surface-available ESC of biochar (∼19%). Together, these results suggest MR-1 accesses the surface-available electron sites of biochar through direct-contact electron transfer. Our results add to the growing body of work demonstrating microbial electron transfer to and from biochar. Understanding the influence of biochar on electron flow and balance within microbial communities is essential to fully evaluating its impact on microbial ecology, nutrient cycling, greenhouse gas production, and contaminant fate when amended or naturally present in the environment.

## Acknowledgements

We thank Jeffrey Gralnick for providing MR-1 and Δ*bfe*, the Delaware Biotechnology Institute (DBI) for shared instrumentation, and Deborah Powell in the Bio-Imaging Center for assistance with SEM.

## References

Afkar E, Reguera G, Schiffer M et al. A novel Geobacteraceae-specific outer membrane protein J (OmpJ) is essential for electron transport to Fe (III) and Mn (IV) oxides in Geobacter sulfurreducens. BMC Microbiol 2005;5:41. 10.1186/1471-2180-5-41.

Barchinger SE, Pirbadian S, Sambles C et al. Regulation of Gene Expression in Shewanella oneidensis MR-1 during Electron Acceptor Limitation and Bacterial Nanowire Formation. Appl Environ Microbiol 2016;82(17):5428–43. 10.1128/AEM.01615-16.

Baron D, LaBelle E, Coursolle D et al. Electrochemical Measurement of Electron Transfer Kinetics by Shewanella oneidensis MR-1*. J Biol Chem 2009;284(42):28865–73. 10.1074/jbc.M109.043455.

Bernardet JF, Bowman J. The prokaryotes: a handbook on the biology of bacteria. Proteobacteria Delta Epsil Subclasses Deep Rooted Bact 2006;7:481–532.

Chen S, Rotaru AE, Shrestha PM et al. Promoting Interspecies Electron Transfer with Biochar. Sci Rep 2014;4(1):5019. 10.1038/srep05019.

Chen Z, Wang Y, Jiang X et al. Dual roles of AQDS as electron shuttles for microbes and dissolved organic matter involved in arsenic and iron mobilization in the arsenic-rich sediment. Sci Total Environ 2017;574:1684–94. 10.1016/j.scitotenv.2016.09.006.

Choi J, Xin D, Chiu PC. A New Climate Impact of Wildfire Chars: Suppression of Biogenic Methane Production Over Repeated Redox Cycles. Environ Sci Technol 2025;59(31):16443–51. 10.1021/acs.est.5c05709.

Coppola AI, Wiedemeier DB, Galy V et al. Global-scale evidence for the refractory nature of riverine black carbon. Nat Geosci 2018;11(8):584–8. 10.1038/s41561-018-0159-8.

Fortune Business Insights. Biochar Market Size, Share, Growth, Trends, Forecast, 2034. 24 Aug. 2026. https://www.fortunebusinessinsights.com/industry-reports/biochar-market-100750 (9 Sept. 2026, date last accessed).

Gorby YA, Yanina S, McLean JS et al. Electrically conductive bacterial nanowires produced by Shewanella oneidensis strain MR-1 and other microorganisms. Proc Natl Acad Sci 2006;103(30):11358–63. 10.1073/pnas.0604517103.

Hong G, Pachter R. Bound Flavin–Cytochrome Model of Extracellular Electron Transfer in Shewanella oneidensis: Analysis by Free Energy Molecular Dynamics Simulations. J Phys Chem B 2016;120(25):5617–24. 10.1021/acs.jpcb.6b03851.

Hunt KA, Flynn JM, Naranjo B et al. Substrate-Level Phosphorylation Is the Primary Source of Energy Conservation during Anaerobic Respiration of Shewanella oneidensis Strain MR-1. J Bacteriol 2010;192(13):3345–51. 10.1128/jb.00090-10.

Jiang X, Hu J, Fitzgerald LA et al. Probing electron transfer mechanisms in Shewanella oneidensis MR-1 using a nanoelectrode platform and single-cell imaging. Proc Natl Acad Sci 2010;107(39):16806–10. 10.1073/pnas.1011699107.

Kappler A, Wuestner ML, Ruecker A et al. Biochar as an Electron Shuttle between Bacteria and Fe(III) Minerals. Environ Sci Technol Lett 2014;1(8):339–44. 10.1021/ez5002209.

Kotloski NJ, Gralnick JA. Flavin Electron Shuttles Dominate Extracellular Electron Transfer by Shewanella oneidensis. mBio 2013;4(1):e00553–12. 10.1128/mBio.00553-12.

Kouzuma A, Kasai T, Hirose A et al. Catabolic and regulatory systems in Shewanella oneidensis MR-1 involved in electricity generation in microbial fuel cells. Front Microbiol 2015;6:609. 10.3389/fmicb.2015.00609.

Li J, Holmes DE, Xu D et al. Fe(III) Oxide Reduction Bypassing Outer-Surface Cytochromes in a Marine Respiratory Anaerobe. Environ Sci Technol 2026;60(20):14601–9. 10.1021/acs.est.6c04532.

Li W, Keffer JL, Singh A et al. Mechanism and capacity of black carbon (biochar) to support microbial growth. Biogeochemistry 2025;168(2):35. 10.1007/s10533-025-01221-y.

Liu X, Ma J, Zhang X et al. Responses of wildfire-induced global black carbon pollution and radiative forcing to climate change. Environ Res Lett 2023;18(11):114004. 10.1088/1748-9326/acff7a.

Loebsack G, Yeung KK-C., Berruti F et al. Adsorption of Organic Pollutants From Wastewater Using Biochar: A Mechanistic Study on Competitive Adsorption Behavior. Water Environ Res 2025;97(8):e70164. 10.1002/wer.70164.

Macheroux P. UV-Visible Spectroscopy as a Tool to Study Flavoproteins. Methods Mol Biol Clifton NJ 1999;131:1–7. 10.1385/1-59259-266-X:1.

Myers CR, Nealson KH. Bacterial manganese reduction and growth with manganese oxide as the sole electron acceptor. Science (New York, N.Y.) 1988;240(4857):1319–21. 10.1126/science.240.4857.1319.

Nevin KP, Lovley DR. Lack of Production of Electron-Shuttling Compounds or Solubilization of Fe(III) during Reduction of Insoluble Fe(III) Oxide by Geobacter metallireducens. Appl Environ Microbiol 2000;66(5):2248–51. 10.1128/AEM.66.5.2248-2251.2000.

Okamoto A, Kalathil S, Deng X et al. Cell-secreted Flavins Bound to Membrane Cytochromes Dictate Electron Transfer Reactions to Surfaces with Diverse Charge and pH. Sci Rep 2014;4(1):5628. 10.1038/srep05628.

Peng B, Chen L, Que C et al. Adsorption of Antibiotics on Graphene and Biochar in Aqueous Solutions Induced by π-π Interactions. Sci Rep 2016;6(1):31920. 10.1038/srep31920.

Pirbadian S, Barchinger SE, Leung KM et al. Shewanella oneidensis MR-1 nanowires are outer membrane and periplasmic extensions of the extracellular electron transport components. Proc Natl Acad Sci 2014;111(35):12883–8. 10.1073/pnas.1410551111.

Qin B, Huang Y, Liu T et al. Dissolved organic matter (DOM) enhances the competitiveness of weak exoelectrogens in a soil electroactive biofilm. Carbon Res 2024;3(1):34. 10.1007/s44246-024-00119-y.

Saquing JM, Yu YH, Chiu PC. Wood-Derived Black Carbon (Biochar) as a Microbial Electron Donor and Acceptor. Environ Sci Technol Lett 2016;3(2):62–6. 10.1021/acs.estlett.5b00354.

Subramanian P, Pirbadian S, El-Naggar MY et al. Ultrastructure of Shewanella oneidensis MR-1 nanowires revealed by electron cryotomography. Proc Natl Acad Sci 2018;115(14):E3246–55. 10.1073/pnas.1718810115.

Sun W, Lin Z, Yu Q et al. Promoting Extracellular Electron Transfer of Shewanella oneidensis MR-1 by Optimizing the Periplasmic Cytochrome c Network. Front Microbiol 2021;12. 10.3389/fmicb.2021.727709.

Tiedje JM. Shewanella—the environmentally versatile genome. Nat Biotechnol 2002;20(11):1093–4. 10.1038/nbt1102-1093.

Tolar JG, Li S, Ajo-Franklin CM. The Differing Roles of Flavins and Quinones in Extracellular Electron Transfer in Lactiplantibacillus plantarum. Appl Environ Microbiol 2023;89(1):e01313–22. 10.1128/aem.01313-22.

Von Canstein H, Ogawa J, Shimizu S et al. Secretion of Flavins by Shewanella Species and Their Role in Extracellular Electron Transfer. Appl Environ Microbiol 2008;74(3):615–23. 10.1128/AEM.01387-07.

Xin D, Li W, Choi J et al. Pyrogenic Black Carbon Suppresses Microbial Methane Production by Serving as a Terminal Electron Acceptor. Environ Sci Technol 2023;57(49):20605–14. 10.1021/acs.est.3c05830.

Xin D, Xian M, Chiu PC. New methods for assessing electron storage capacity and redox reversibility of biochar. Chemosphere 2019;215:827–34. 10.1016/j.chemosphere.2018.10.080.

Xing J, Dong W, Liang N et al. Sorption of organic contaminants by biochars with multiple porous structures: Experiments and molecular dynamics simulations mediated by three-dimensional models. J Hazard Mater 2023;458:131953. 10.1016/j.jhazmat.2023.131953.

Yang Y, Ding Y, Hu Y et al. Enhancing Bidirectional Electron Transfer of Shewanella oneidensis by a Synthetic Flavin Pathway. ACS Synth Biol 2015;4(7):815–23. 10.1021/sb500331x.

Yang Z, Sun T, Kappler A et al. Biochar facilitates ferrihydrite reduction by Shewanella oneidensis MR-1 through stimulating the secretion of extracellular polymeric substances. Sci Total Environ 2022;848:157560. 10.1016/j.scitotenv.2022.157560.

Yu L, Wang Y, Yuan Y et al. Biochar as Electron Acceptor for Microbial Extracellular Respiration. Geomicrobiol J 2016;33(6):530–6. 10.1080/01490451.2015.1062060.

