## Supplemental file for "*Shewanella oneidensis* grows on biochar as an electron acceptor via direct contact without flavins"

*Corresponding authors:

### Summary (4 pages total, 1 table, 3 figures)

This document contains (1) Experimental growth data with WT MR-1 and Δ*bfe* MR-1 to confirm similar anaerobic growth on a soluble electron acceptor. (2) Electron donation curves from redox titration measurements of post-growth and control biochar electron storage capacity. (3) Additional images of WT MR-1 and Δ*bfe* MR-1 on biochar. (4) Data table with electron donor consumption and electron acceptor usage numbers.

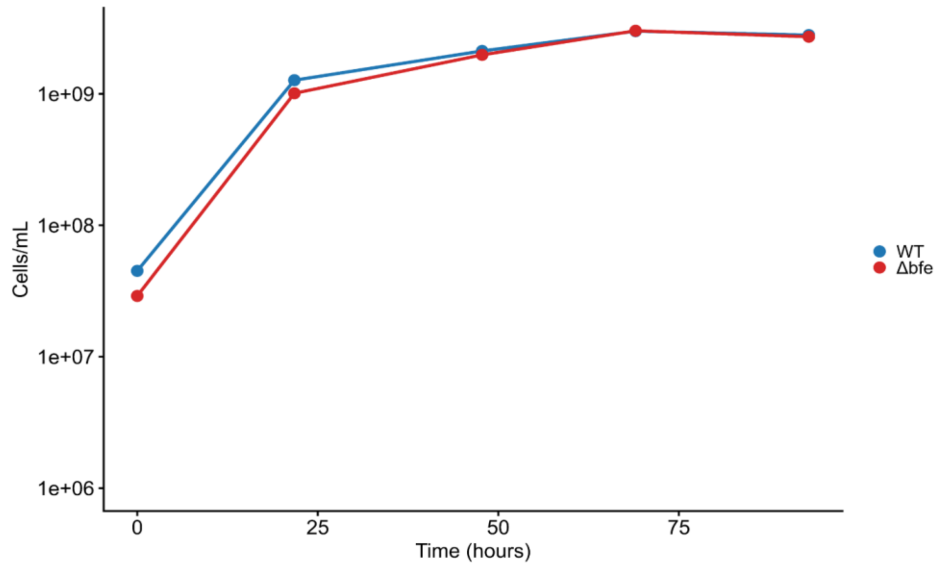


**Figure S1.** Wildtype MR-1 and Δ*bfe* MR-1 growth anaerobically using lactate as an electron donor and fumarate as an electron acceptor. Both strains show similar growth, indicating no growth defects caused by removal of the flavin exporter gene.

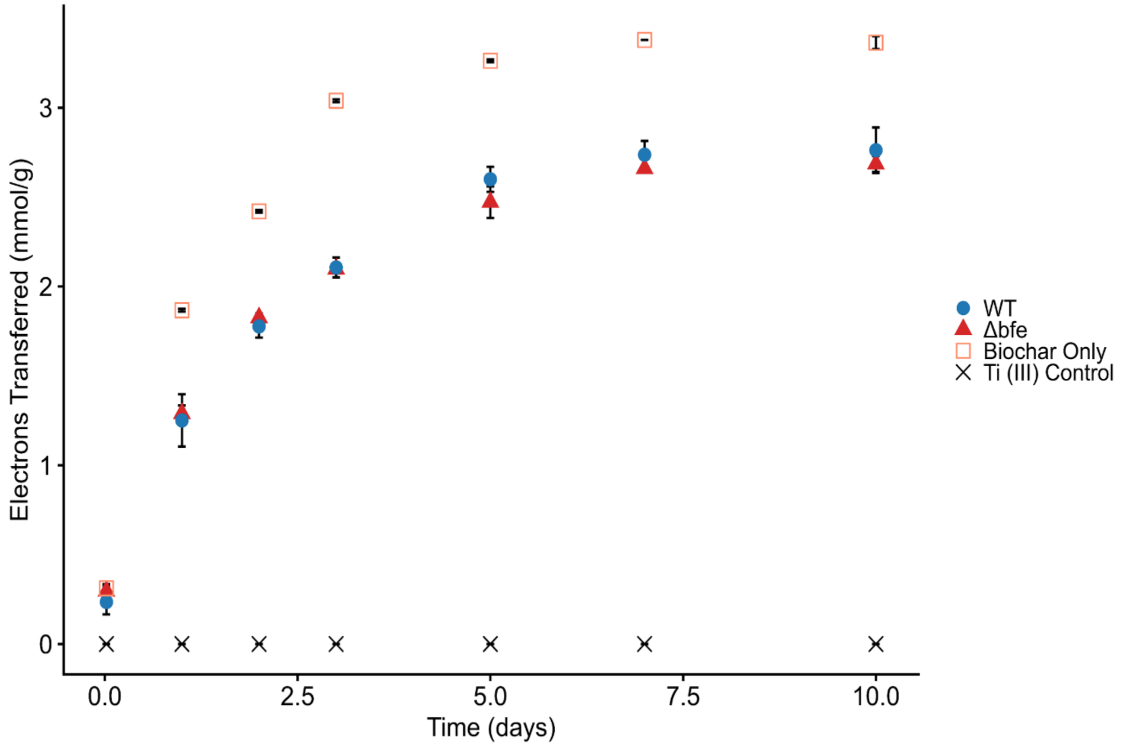


**Figure S2:** Chemical redox titration results for WT-reduced biochar, Δ*bfe*-reduced biochar, and biochar in media with no cells harvested after the growth experiment. Error bars represent one standard deviation based on triplicates.

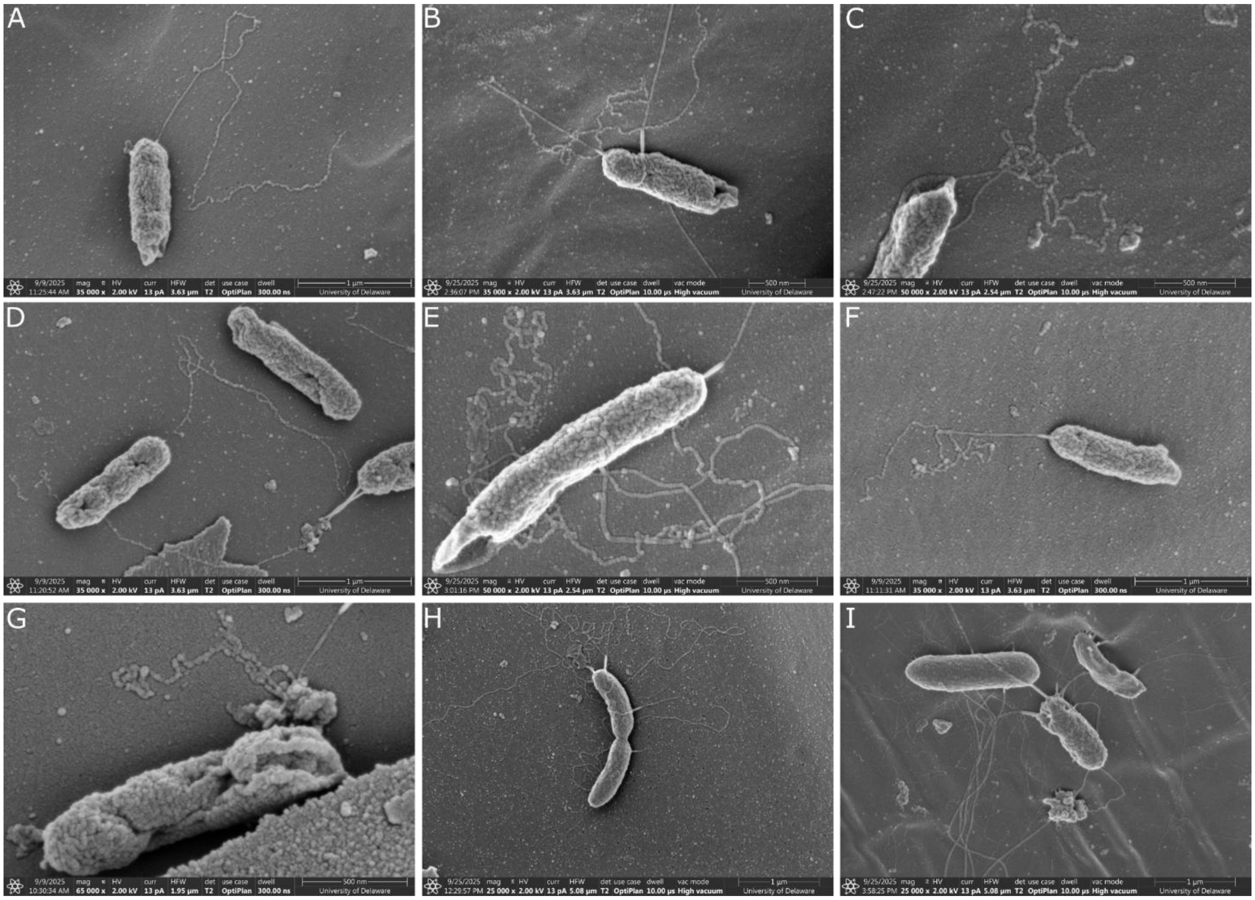


**Figure S3**. Additional images of WT (A-F) and Δ*bfe* (G-I) showing nanowire contacts between cells and biochar.

**Table S1.** Electron donor (lactate) consumption and electron acceptor (biochar) capacity measurements.

| Strain | Lactate Consumed (mM) | Electron Consumption Calculated from Lactate (mmol) | Total Biochar Capacity Remaining (mmol/g) | Electron Deposition to Biochar (mmol) |
|---|---|---|---|---|
| **Biochar only** | - | - | 3.4 ± 0.04 | - |
| **WT** | 7.4 ± 1.2 | 1.5 ± 0.23 | 2.8 ± 0.13 | 0.52 ± 0.11 |
| **WT (no e-acceptor)** | 2.4 ± 3.6 | 0.48 ± 0.73 | - | - |
| **Δ*bfe*** | 6.4 ± 0.90 | 1.3 ± 0.18 | 2.7 ± 0.04 | 0.66 ± 0.05 |
| **Δ*bfe* (no e-acceptor)** | 0.57 ± 1.9 | 0.11 ± 0.38 | - | - |
